# The carrier proteome limit should be reassessed for each mass analyzer architecture

**DOI:** 10.1101/2024.03.08.584130

**Authors:** Benjamin C. Orsburn

**Author notes:** **Corresponding: Benjamin C. Orsburn**, The Johns Hopkins University School of Medicine, Department of Pharmacology and Molecular Sciences, 725 North Wolfe Street, Baltimore, MD 21205, **Email:**. **Author Contributions:** Conceptualization: BCO Methodology: BCO Investigation: BCO Funding acquisition: BCO Writing – original draft: BCO Writing – review & editing: BCO. **Competing Interest Statement:** I have no competing interests to disclose.

## Abstract

A clever utilization of classic proteomics reagents now allows the effective amplification of peptide sequencing potential in shotgun proteomics. The application of this method has helped usher in the exciting new field of single cell proteomics. While it was easy to first think that the discovery of Budnik et al., was finally the answer for protein PCR, limitations were carefully described by the authors and others. A study by Cheung et al., systematically identified the consequences of higher concentration carrier proteomes and defined the “carrier proteome limit”. While this work has been replicated by others, every analysis published to date has used a variation of the same mass analyzer. When the same analysis is performed on alternative instruments, these limits appear to be very different and attributable to defined characteristics of each mass analyzer. Specifically, in mass analyzers with higher relative intrascan linear dynamic range, increased carrier channels appear far less detrimental to quantitative accuracy. As such, we may be limiting the power of isobaric peptide signal “amplification” by restricting ourselves to traditional mass analyzer options.

## Main

Leveraging the cumulative signal of isobarically tagged peptides in multiplexed proteomics has helped drive the emerging new field of single cell proteomics by mass spectrometry (SCP). While isobaric tagging reagents have been in use for decades, Budnik *et al*., seems to have been the first description of the use of higher concentration “carrier proteomes” to effectively amplify peptide sequencing potential.^1^ Protein biochemists have long dreamed of finding an equivalent to the oligonucleotide polymerase chain reaction (PCR). However, an analysis by Cheung *et al*., appeared in Nature Methods in short order to temper these expectations by defining the upper limits, and consequences of approaching or exceeding those limits, in a carrier proteome enabled experiment.^2^ These researchers found that as the carrier proteome increased, quantitative accuracy tended to decrease. While the math differs a little, these results were verified in multiple successive follow-up studies.^3,4^

However, every one of these studies utilized some version of hybrid Orbitrap mass analyzer. This isn’t surprising, as hybrid Orbitrap instruments have been the most utilized instrument configuration for LCMS based proteomics for over a decade. Orbitraps have high mass accuracy and resolution and have steadily improved as proteomic instruments since their introduction.^5^ However, a concrete limitation of any ion trap is that charged ions influence one another, and that too many ions in a trap will invariably lead to measurement errors known as space charging effects. Orbitraps are no exception, and neither are the simpler accessory ion traps that aid in their operation deep within the instruments.^5^ One challenge of having a finite capacity of ions is that you also have a finite intrascan linear dynamic range (ILDR), which is typically defined as the highest and lowest number of ions within a single mass spectrum where a linear response is maintained. While numbers can vary between different configurations, about 1 x 10^6^ ion charges can fit within an Orbitrap without bad things happening, and when you distribute these charges between low and high abundance ions, you’ll find that you’ve got an ILDR of about 2 orders of magnitude.^6^ It does not appear coincidental, therefore, that every study on carrier proteome limits has found detrimental effects when a carrier is more than 2 orders higher than the single cell peptide concentration.^2–4^

In contrast, time of flight (TOF) mass analyzers classically do not perform any steps where ions are held in space or otherwise forced to interact with one another in a confined space. The ILDR of these analyzers have long been known to be significantly higher than ion trapping instruments for these reasons.^6,7^ In support of the ILDR directly influencing the carrier proteome limits, we recently reported less detrimental carrier channel effects on a trapped ion mobility TOF (TIMSTOF) than observed for Orbitrap hardware.^8^ These results were supported by an independent study which observed no detriment to using a 200x carrier channel on a similar instrument.^9^

To further evaluate the link between the ILDR and carrier proteome limits, I repeated our carrier limit assessment on a recently described TOF instrument with a reported ILDR of 5 orders of magnitude, which is higher than any instrument reported to date for a carrier proteome enhanced experiment.^10^

Similar to previous analyses, I re-utilized a complex peptide digest mixture containing a constant complex background mixed with peptides at known ratios.^2,3^ While all human proteins detected should be at a 1:1 ratio in every multiplex channel, all *E*.*coli* peptides detected should be at different known concentrations in each channel. The only factor that changes is the amount of the carrier proteome channel that is added in the final mixture of sample. As reported by Cheung *et al*., and others our previous analysis of this same standard on our Orbitrap instrument shows a rapid compression of the known 5:1 ratio for the *E*.*coli* proteins. However, when analyzing these same samples on a TIMSTOF second generation instrument we observed a much smaller degree of ratio compression. A summary figure from one of these analyses in the known vs observed values for the *E*.*coli* protein TNAa is presented as **Figure 1A**.^8^ In addition, as shown in **Figure 1B**, the higher ILDR TOF instrument has even less apparent alteration in the observed ratios of this protein even up to a carrier proteome 441 fold higher than the other channels. While it can be noted that the ZenoTOF system experiences considerable ratio compression, this is a struggle for all mass analyzers, particularly those where ion isolation is exclusively performed by quadrupole isolation.^11,12^ The metric of note here should be that on a second mass analyzer architecture, the carrier proteome limits appear entirely different than the multiple existing reports for this effect on Orbitrap hardware.

**Figure 1.**
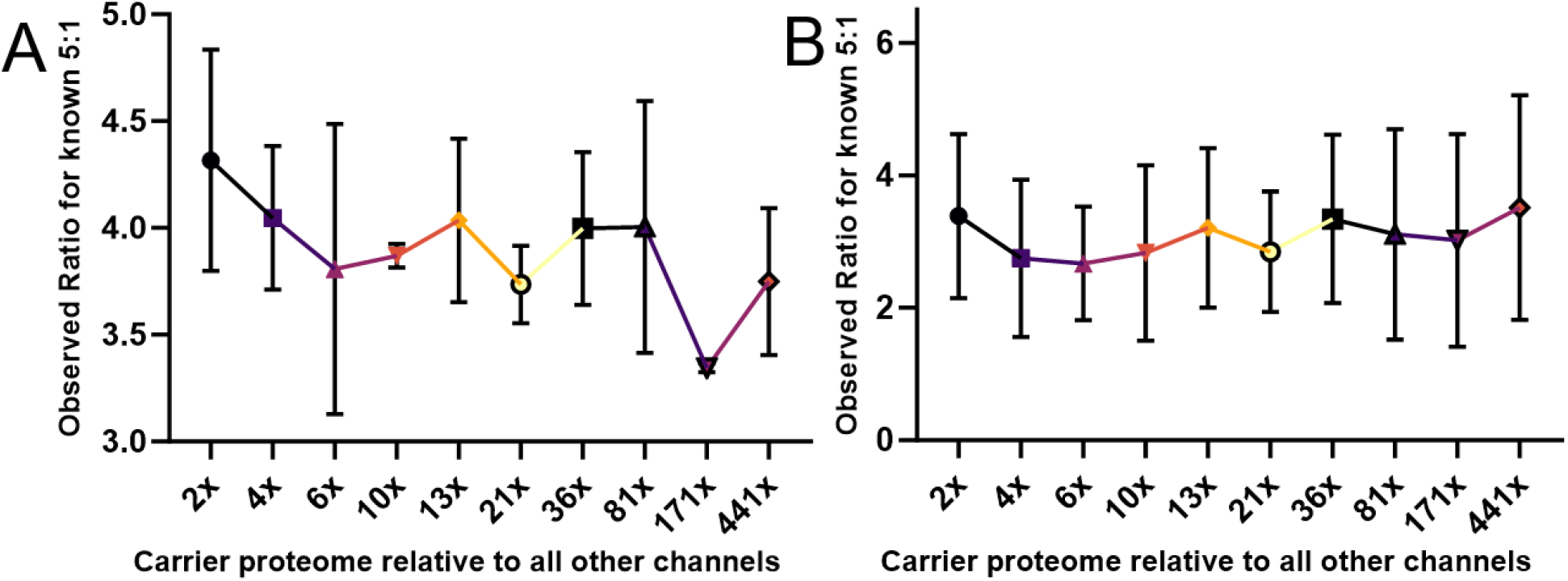
The effects of increasing the carrier proteome channel in a complex 2-proteome digest on TOF instruments for the TNAa protein which has a known 5:1 ratio between channels. **A**. Previously published data for this standard on a TIMSTOF system.^8^ **B**. The reanalysis of these samples presented here on a ZenoTOF system. Mean ratio for each observed peptide and standard deviation are displayed. Source data are provided.

In conclusion, these results suggest that the carrier proteome limit should be carefully evaluated for each mass analyzer architecture. It is tempting to think that the true potential of the isobaric peptide amplification trick described by Budnik *et al*.,^1^ may actually be closer to the power of PCR than multiple studies have indicated.

## Methods

The sample used to assess the carrier channel effect was previously described. Briefly, a K562 human peptide digest standard (Promega) and *E*.*coli* peptide standard (Waters) were resuspended in 100mM TEAB (SimpliFi) and aliquots of 10 micrograms of each were labeled with the TMTPro reagents, 126, 127n, 128n, 129n, 130n, 131n, 132n, 133n and 134 and the reactions quenched with hydroxylamine according to vendor instructions. Labeled peptides were combined to maintain and equal concentration of K562 digest in each lane by combining 2 micrograms of each. *E*.*coli* peptides were diluted to 10:5:1 in TEAB and combined with the respective lanes to result in a repeating E.coli concentration of 20% and 10% of the first channel. The combined peptides were diluted to 100ng/μL total concentration and analyzed on a SCIEX 7600 ZenoTOF instrument coupled to a Waters H Class microflow UHPLC. A 60 minute gradient was used at a flow rate of 5 μL/min using a Waters BEH C-18 NanoEaze column 300 μm x 150mm with 1.8 μm particles. The gradient ran from 95% buffer A (0.1% formic acid in LCMS grade water) to 35% buffer B (0.1% formic acid in LCMS grade acetonitrile) in 45 min before ramping to 90% B in 10 min and returning to baseline conditions. A top 10 data dependent method was used for all analysis. The resulting output files were converted to MGF using the ProteoWizard software and the MGF files were processed in Proteome Discoverer 2.4SP1 as previously described.^8^

## Data Availability

All vendor .wiff files, converted data and processed results are publicly available through the ProteomeXchange partner repository as accession PXD049286. TIMSTOF carrier proteome data reanalyzed in this study was previously published as part of accession PXD028710.

